# Architectural Confinement and Seasonal Forcing Shape Cross-Domain Pathogen Assemblages in Complex Built Environments

**DOI:** 10.64898/2026.05.26.727965

**Authors:** Shijiao Qi, Shurui Zhang, Yixin Hu, Mingjie Sun, Zichun Xing, Shanshan Bao, Yuyang Song, Li Sun, Xinzhao Tong

## Abstract

Airborne microbial pathogens in built environments (BEs) may pose potential health threats, yet the ecological mechanisms governing their assembly and persistence across interconnected architectural spaces are less studied. Here, we conducted a year-long, multi-spatial investigation across a university campus complex, sampling the exhaust outlets of three indoor environments and the adjacent outdoor inlets. By integrating the 16S rRNA gene and ITS sequencing from 437 paired bacterial–fungal samples, we characterized the spatiotemporal dynamics of airborne opportunistic pathogen-containing genera. Our study showed that spatial filtering emerged as the dominant determinant of pathogen community structure, with confined elevator environments serving as reservoirs of pathogens with limited but continuous microbial influx from surrounding spaces. Superimposed on this spatial baseline, bacterial and fungal pathogens displayed distinct seasonal dynamics. Opportunistic fungal pathogens such as *Fusarium* peaked during autumn and winter, possibly driven by enhanced aerosol persistence and dispersal under cooler, drier conditions. In contrast, bacterial pathogens exhibited greater temporal resilience, with key taxa such as *Listeria* transcribed actively during winter, predisposing them to increased relative abundance during spring. Cross-domain ecological networks further revealed the dynamic interactions among pathogens, centered on the skeletal structure mediated by the keystone fungal pathogen *Aspergillus*, suggesting that coordinated microbial interactions reinforced pathogen persistence across seasons. Together, our findings support a unified ecological framework in which pathogen dynamics within BEs arise from the coupled effects of spatial filtering, climatic modulation, and biotic interactions. These results provide a foundation for transitioning from static environmental control toward predictive pathogen management in BEs.

**Importance:** BEs are the primary settings of human microbial exposure, yet the ecological principles governing the persistence of airborne pathogens across interconnected indoor spaces remain poorly resolved. Most previous indoor studies have focused on isolated environments or single microbial groups, limiting our understanding of how pathogens are maintained and redistributed within complex architectural systems. By integrating bacterial and fungal community dynamics across spatial environments over four seasons, this year-long study demonstrates that the ecology of airborne pathogens is not static, but is instead dictated by a complex interplay of spatial, climatic, and biological forces. Our findings identify enclosed, high-transit elevator spaces as critical hotspots for pathogen accumulation and highlight the role of seasonal ecological reorganization in driving airborne health risks. More broadly, this work establishes a systems-level ecological framework for understanding airborne pathogen dynamics in BEs and provides a conceptual basis for developing more adaptive strategies for indoor microbial risk management.

## Introduction

With rapid urbanization in the 21st century, increasingly complex built environments (BEs) have become essential to support dense populations and expanding infrastructure systems. As a consequence, modern human activities are overwhelmingly confined to indoor spaces, including residential, commercial, and transportation settings (1). Despite routine cleaning and disinfection, these environments are far from sterile. Instead, they harbor diverse and dynamic airborne microbial communities composed of bacteria, fungi, and other microorganisms. Among these, exposure to a subset of opportunistic pathogens may pose substantial health risks through inhalation and deposition in the respiratory tract. Common airborne bacterial genera such as *Legionella*, *Mycobacterium*, *Staphylococcus*, and *Pseudomonas*, and fungal taxa including *Aspergillus*, *Cladosporium*, *Penicillium*, and *Alternaria*, are frequently detected in BEs and have been associated with respiratory infections, allergic diseases, and asthma exacerbation (2).

The presence of these microorganisms is not random but governed by ecological processes that regulate their composition and persistence. Airborne microbial communities in BEs are shaped by the combined effects of environmental selection and dispersal dynamics (2). Environmental and building factors, including temperature, relative humidity, ventilation rate, light availability, and building materials, act as selective filters that determine microbial survival, growth, and activity (3). At the same time, dispersal processes mediated by airflow, human movement, and architectural connectivity influence microbial transmission across spaces (2). Together, these processes generate spatially heterogeneous microbial communities within buildings.

Such heterogeneity is particularly evident across distinct architectural environments. Underground spaces are typically characterized by elevated humidity, more stable temperatures, limited light, and restricted ventilation, which may serve as reservoirs for humidity-associated pathogens, including toxigenic fungi, that can be redistributed via ventilation-mediated transport (4). In contrast, aboveground indoor environments are more strongly influenced by outdoor air exchange and diurnal environmental fluctuations. Confined microenvironments, such as elevators, represent highly connected nodes with heavy human traffic, frequent surface contact, and limited air circulation, creating unique ecological conditions for microbial accumulation and redistribution. These differences not only shape microbial diversity but also influence the persistence and transmission potential of pathogens (5). Therefore, enclosed and high-traffic spaces may facilitate microbial homogenization and act as hubs for pathogen accumulation and dispersal, thereby increasing the likelihood of aerosol transmission through shared airflows and ventilation systems (5).

Beyond spatial heterogeneity, temporal variability further modulates microbial dynamics. Seasonal climatic fluctuations alter environmental conditions that govern microbial growth and dispersal (6), an effect that is especially pronounced in subtropical monsoon regions where temperature and humidity vary substantially across seasons. The Yangtze River Delta, one of the most densely populated and rapidly urbanized regions in China, exemplifies this climatic regime. A defining feature of this region is the annual plum rain season, characterized by persistent precipitation, elevated humidity, and warm temperatures during June and July (7). These conditions sustain high indoor moisture levels and create favorable environments for microbial proliferation, particularly for fungi and moisture-associated bacterial taxa (2). Although climatic influences on indoor microbiomes have been widely recognized, the interactive effects of spatial heterogeneity and seasonal climatic variability across vertically connected public BEs remain insufficiently characterized.

In this study, we conducted a year-long investigation (September 2023–August 2024) on a university campus in the Yangtze River Delta, serving as a model system for complex public buildings with interconnected, functionally distinct compartments. We continuously monitored temperature and relative humidity across four spatial settings: aboveground indoor spaces, underground facilities, elevators, and outdoor environments. Concurrently, bacterial and fungal communities were characterized using 16S rRNA gene (16S rDNA), 16S rRNA, and ITS sequencing of swab samples collected from the outlets and inlets of the ventilation systems. A total of 437 paired bacterial–fungal samples were obtained, enabling a comprehensive assessment of microbial diversity and composition across spatial and temporal gradients.

This study aims to (i) characterize the spatiotemporal dynamics of bacterial and fungal communities across distinct spatial environments, (ii) identify key environmental drivers shaping microbial diversity, composition, and metabolic activity, (iii) evaluate the distribution and dispersal potential of pathogenic microorganisms across interconnected indoor spaces, and (iv) explore potential interactions between bacterial and fungal taxa, with particular emphasis on co-occurrence patterns among microbes with pathogenic potential. By integrating microbial ecology with architectural and climatic factors, understanding these dynamics is essential for designing healthier BEs and mitigating risks of airborne pathogen transmissions.

## Materials and Methods

### Study Design, Sample Collection, and Sequencing

Full methodological details are provided in the supplementary material (**Text S1**). This study was conducted in the Science Buildings of Xi’an Jiaotong-Liverpool University (Suzhou, China), a complex featuring extensive interconnected subterranean and aboveground infrastructure. Swab samples were collected biweekly over an annual cycle (September 2023 to August 2024) from the ventilation exhaust surfaces of three interconnected indoor environments (aboveground spaces, underground facilities, and enclosed elevators), as well as from adjacent outdoor air inlets (**Figure S1**). Ambient temperature and relative humidity were recorded at each sampling event.

Genomic DNA was extracted from all samples using the QIAGEN DNeasy PowerSoil Pro Kit. Amplicon sequencing of the bacterial 16S rRNA gene (V4 region) and fungal ITS1 region was executed on the Illumina MiSeq platform, yielding 437 high-quality, matched bacterial–fungal sample groups after quality filtering. Concurrently, to profile active transcriptional states during two highly contrasting seasons (winter vs. the plum rain summer), total RNA was extracted from 36 parallel underground samples, reverse-transcribed to cDNA, and subjected to bacterial 16S rRNA amplicon sequencing.

### Bioinformatic Processing and Statistical Analyses

Raw sequences were processed via the QIIME2 pipeline (v2024.2) (9), utilizing DADA2 (10) for denoising and generating amplicon sequence variants (ASVs). Bacterial and fungal ASVs were taxonomically assigned against the Greengenes2 (v2024.09) (12) and UNITE (v9.0) (13) databases, respectively. Pathogen-associated genera were inferred by matching our taxonomy tables against genus-level information from the Global Catalog of Pathogens (gcPathogen) database (17), restricting the criteria to species classified as Biosafety Level 2 or higher.

Following sample rarefaction to even depths, alpha-diversity was evaluated via the Shannon index and observed richness. Long-term and seasonal temporal trajectories in alpha-diversity were modeled using generalized additive models (GAMs) via the R package “mgcv” (v1.9-4). For beta-diversity, community composition patterns were evaluated using abundance-based Bray–Curtis dissimilarity matrices and visualized through Principal Coordinates Analysis (PCoA). Permutational multivariate analysis of variance (PERMANOVA) and analysis of multivariate dispersions (PERMDISP) in the R package “vegan” (v2.7.3) were applied to assess compositional significance and dispersion homogeneity across spatial and temporal groups.

The effects of environmental parameters (temperature, relative humidity, and their interactions) on pathogen abundance and prevalence were modeled using linear mixed-effects and logistic regressions in MaAsLin3 (v0.99.16) (19). Bacterial relative transcriptional activity was estimated from the rRNA/rDNA ratio of corresponding ASVs, and metabolically active taxa were defined as those with a ratio >1 (8). Cross-domain community concordance was evaluated using Procrustes analysis (protest function) and Mantel tests on the respective Bray–Curtis dissimilarity matrices, implemented in the R package “vegan”. Cross-domain microbial co-occurrence networks were constructed using the SPIEC-EASI algorithm (22) to resolve associations between bacteria and fungi. Finally, bidirectional microbial dispersal routes and migration rates (*m*) among four spatial groups were simulated using the Sloan neutral community model (NCM) (v2015.10) (20).

## Results

### Spatiotemporal Dynamics of Bacterial and Fungal Communities

At the community level, bacterial assemblages were consistently dominated by four major classes throughout all seasons: Bacilli, Actinobacteria, Alphaproteobacteria, and Gammaproteobacteria (**Figure 1a**). However, their temporal dynamics showed a spatially dependent pattern. In elevators, these taxa maintained relatively stable abundances across seasons, with Gammaproteobacteria and Actinobacteria particularly enriched. In contrast, aboveground and underground environments exhibited pronounced seasonal variation: Bacilli gradually accumulated from autumn to summer in underground spaces, whereas in aboveground settings, they peaked in spring before declining. Compared to the bacterial community, the fungal counterpart displayed markedly different, less diverse class-level patterns. Dothideomycetes dominated outdoor, aboveground, and underground environments, but were disproportionately depleted in elevators (**Figure 1b**). Furthermore, Eurotiomycetes were less abundant in aboveground settings than the other spatial types, and Sordariomycetes showed consistent winter peaks across all spatial types.

**Figure 1.**
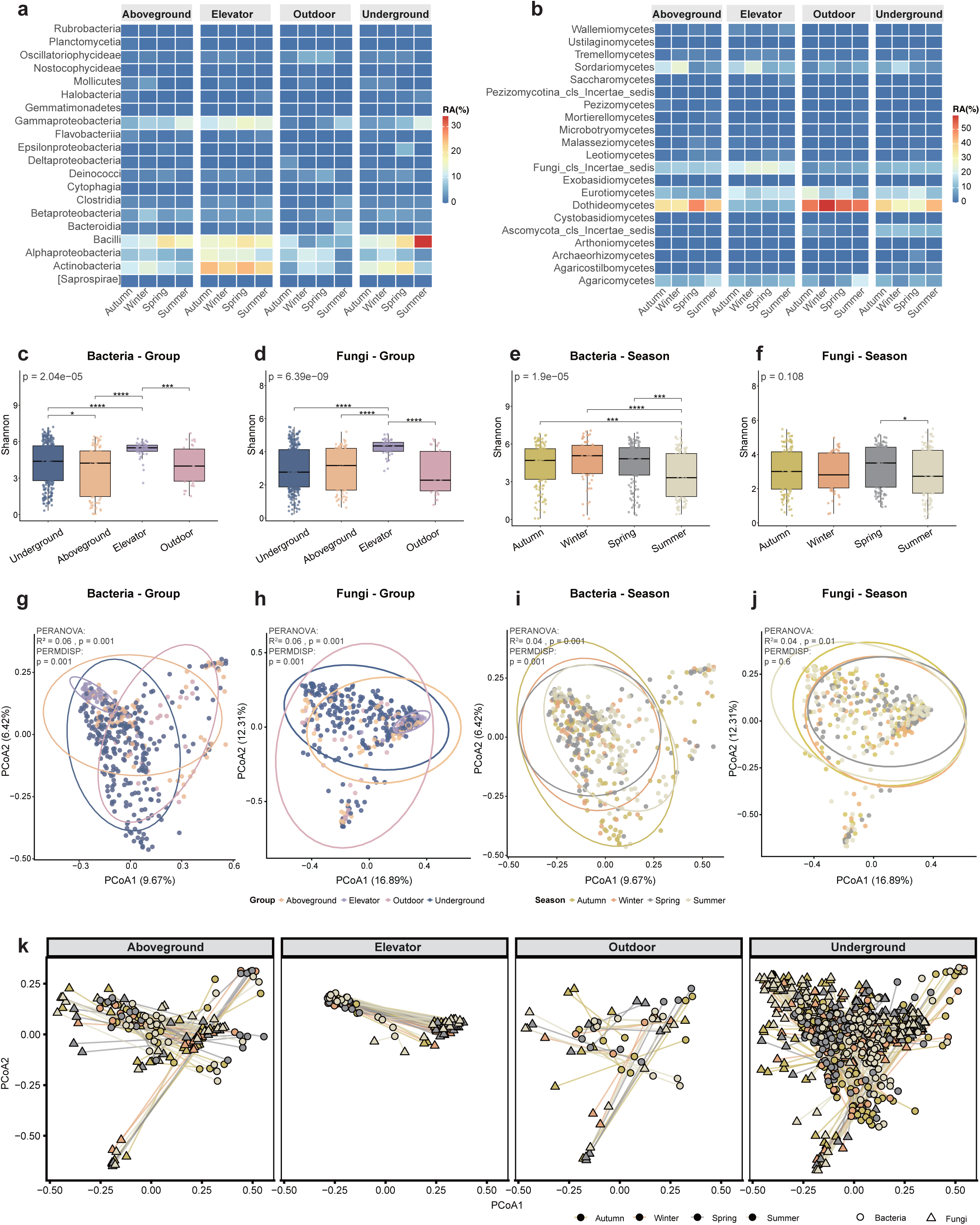
Spatial and seasonal structuring of airborne microbial communities. **a,b)** Relative abundances of the top 20 bacterial **(a)** and fungal **(b)** classes across all samples. **c, d)** Shannon diversity of bacterial **(c)** and fungal **(d)** communities across four spatial environments. **e, f)** Shannon diversity of bacterial **(e)** and fungal **(f)** communities across four seasons. **g, h)** Principal coordinates analysis (PCoA) of bacterial **(g)** and fungal **(h)** communities across spatial environments, using Bray–Curtis dissimilarity for both domains. Ellipses represent 95% confidence intervals for group centroids. **i, j)** PCoA of bacterial **(i)** and fungal **(j)** communities across seasons using the same dissimilarity metrics as indicated above. **k)** Procrustes-based coupling between bacterial and fungal community structures along spatial gradients, indicating coordinated shifts across environments.

The overall Shannon diversity of bacterial and fungal communities was positively correlated (Spearman’s ρ = 0.681, *P* = 5.65 × 10^-34^), suggesting partially shared ecological drivers across time and space. When spatial setting was considered, significant differences in alpha diversity were detected for both domains (Kruskal-Wallis test, *P*_bacteria_ = 2.04 × 10^-5^, *P*_fungi_= 6.39 × 10^-9^), following a parallel gradient in which elevator environments consistently harbored the highest diversity, while outdoor settings exhibited the lowest diversity (**Figures 1c and 1d**). Conversely, temporal dynamics diverged between domains. Bacterial diversity differed significantly across seasons (*P* = 1.90 × 10^-5^), peaking in winter and declining in summer, whereas fungal diversity showed no statistically significant seasonal changes (*P* = 0.108), though there was a tendency toward higher values in spring (**Figures 1e and 1f**). Overall, the alpha-diversity analyses revealed consistent spatial gradients, alongside divergent temporal responses in both bacterial and fungal communities.

To quantify the drivers of microbial alpha diversity, GAMs were fitted separately for bacteria and fungi. The models explained 19.8% and 16.5% of the deviance in bacterial and fungal richness, respectively (bacteria: *adj. R²* = 0.179; fungi: *adj. R²* = 0.146, **Figures S2a and S2b**). Across both domains, spatial environment type emerged as the dominant and statistically significant determinant of microbial richness in both domains, accounting for the largest single reduction in explained variance upon removal (*ΔR²* = 0.077 for bacteria; *ΔR²* = 0.096 for fungi), substantially exceeding the contributions of all temporal and climatic components combined (*ΔR²* ≤ 0.021). Regarding temporal dynamics, no significant global long-term trend was detected for bacteria (edf = 1.00, *P* > 0.05) and fungi (edf = 1.00, *P* > 0.05), and space-specific trend deviations were also non-significant for bacteria across all spatial types (all *P* > 0.05, **Figure S2a**). However, fungal richness in underground environments exhibited a significant environment-specific nonlinear temporal trend (edf = 3.55, *F* = 3.62, *P* = 2.37 × 10^-4^, **Figure S2b**), indicating complex, non-monotonic temporal dynamics in fungal communities. This underground-specific temporal deviation contributed the most to explained variance among all smooth components (*ΔR²* = 0.023 for fungi). Seasonal dynamics were similarly space-specific. No significant global seasonal pattern was detected in either domain (**Figure S2c and S2d)**. Nevertheless, bacterial richness in underground environments showed a significant seasonal deviation from the global pattern (edf = 3.20, *F* = 2.45, *P* = 5.37 × 10^-4^, **Figure S2c**), reflecting complex within-year cyclical fluctuations unique to ventilation surface environments (*ΔR²* = 0.003). No seasonal signals were detected for fungi in any spatial environment (all *P* > 0.05, **Figure S2d**), indicating that fungal richness may not follow predictable within-year cycles in any of the sampled environments.

Beta-diversity analyses similarly revealed significant spatial structuring differences in both bacterial and fungal communities (PERMANOVA *R^2^_bacteria_* = 0.06, *P*_bacteria_ = 0.001, *R^2^_fungi_* = 0.06, *P*_fungi_ = 0.001, **Figures 1g and 1h**). However, community dispersion differed significantly among spatial types in both domains (PERMDISP *P* = 0.001 for both domain), suggesting that compositional variation could be driven by heterogeneous dispersion across different spaces. This was primarily attributable to the pronounced clustering of elevator samples relative to the broader scatter observed elsewhere, indicating strong environmental filtering within enclosed, human-managed habitats. Seasonal effect on fungal community composition was weak but significant (PERMANOVA *R*^2^ = 0.04, *P* = 0.001; PERMDISP *P* = 0.6, **Figure 1j**), whereas bacterial communities primarily varied in dispersion rather than centroid position (PERMANOVA *R*^2^ = 0.04, *P* = 0.001; PERMDISP *P* = 0.001, **Figure 1i**), highlighting distinct modes of temporal variability between the two domains.

Bacterial–fungal coupling closely tracked these spatial gradients (**Figure 1k**). Across all samples, the two domains exhibited strong concordance (Procrustes *r* = 0.84, *P* = 0.001; Mantel *r* = 0.52, *P* = 0.001). Although significant alignment was detected within all individual spatial types (all *P* = 0.001), coupling strength decreased progressively from elevators (Procrustes *r* = 0.982/0.186, *SS* = 0.642; Mantel *r* = 0.501) to aboveground (Procrustes *r* = 0.907/0.420, *SS* = 0.266; Mantel *r* = 0.590), underground (Procrustes *r* = 0.782/0.623, *SS* = 0.311; Mantel *r* = 0.505), and outdoor environments (Procrustes *r* = 0.709/0.705, *SS* = 0.239; Mantel *r* = 0.731). Procrustes residuals differed significantly among spatial types (ANOVA, *F* = 13.12, *P* = 3.26 × 10^-8^), with smaller residuals in enclosed environments reflecting tighter cross-domain congruence compared to the greater variability observed outdoors (**Figure S3**).

Taken together, these findings reveal that spatial context overwhelmingly governs the dynamics of both bacterial and fungal communities at a localized scale. Enclosed environments, such as elevators, exhibited a distinct microbial profile: reduced beta-diversity suggests that physical shielding maintains stable community composition, while elevated alpha-diversity indicates a consistent influx of taxa from external sources, such as human occupants, jointly resulting in tightly clustered yet taxonomically rich assemblages with strong cross-domain coupling. While microclimatic variables and seasonal effects also contribute to community assembly, they tend to act as secondary modulators, ultimately constrained by the dominant spatial filtering.

### Spatiotemporal Structure of Pathogenic Microbial Assemblages

Building upon the whole-community patterns, we next examined whether pathogen-associated taxa recapitulate these spatiotemporal rules or exhibit distinct ecological dynamics. Our results showed that fungal pathogens were, on average, consistently more abundant than their bacterial counterparts across all spatial environments; the former peaked in abundance during autumn and winter, whereas the latter reached maximum abundance in spring and summer. When accounting for spatial groups, pathogenic assemblages in both domains followed a distinct, matching gradient of relative abundance: aboveground > elevators > underground > outdoor (**Figure 2a** and **2b**). The alpha diversity of pathogenic bacterial and fungal communities largely mirrored the patterns of the total microbial community (*P*_bacteria_ = 1.4 × 10^-4^, *P*_fungi_ = 3.23 × 10^-10^, with elevators consistently harboring the highest pathogen diversity across all seasons (**Figures 2c** and **2d**). This suggests that rather than acting as a restrictive ecological filter, enclosed transit hubs such as elevators may function as reservoirs of diverse pathogens. Furthermore, the seasonal dynamics of pathogenic diversity generally tracked those of the total community (**Figures S4a** and **S4b**). However, this temporal signature was significantly intensified in the fungal pathogenic subset relative to the total mycobiome, characterized by a shape decline in winter and a prominent peak in spring (**Figure S4b**).

**Figure 2.**
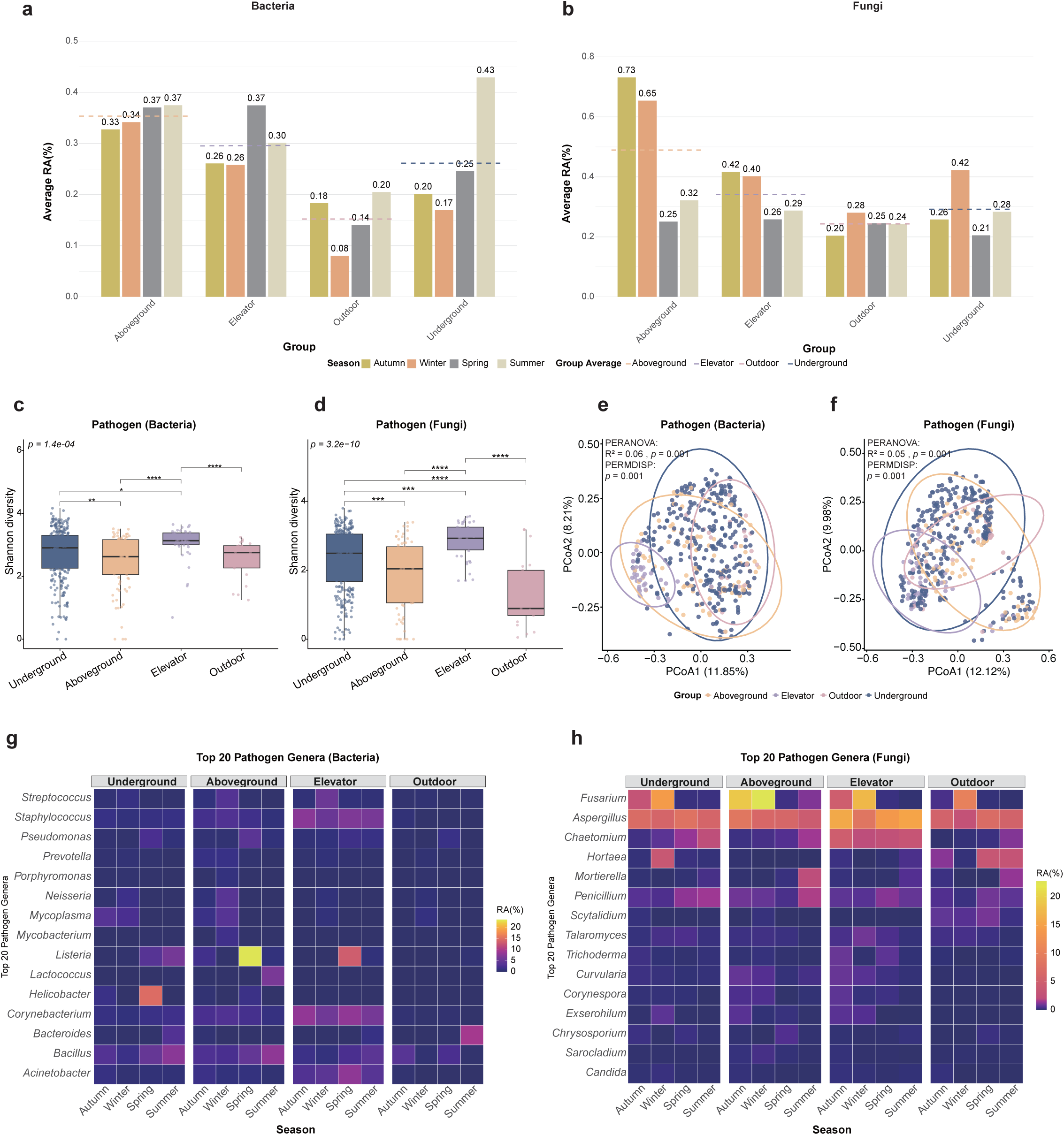
Spatial and seasonal variation in airborne pathogen diversity and composition. **a,b)** Summed relative abundance of pathogen-associated bacterial **(a)** and fungal **(b)** genera across spatial environments and seasons. Bars represent seasonal means within each spatial category; dashed lines indicate overall means per environment. **c, d)** Shannon diversity of bacterial **(c)** and fungal **(d)** pathogen communities across four spatial environments. **e, f)** Principal coordinates analysis (PCoA) of bacterial **(e)** and fungal **(f)** pathogen-associated communities across spatial environments, with Bray–Curtis dissimilarity metric applied for both domains. **g, h)** Heatmaps showing the relative abundance of the 20 most prevalent bacterial **(g)** and fungal **(h)** pathogenic genera across spatial environments and seasons.

Beta-diversity analyses similarly revealed significant spatial differentiation in both bacterial and fungal pathogenic communities (**Figures 2c** and **2d**, PERMANOVA *R^2^_bacteria_* = 0.06, *P*_bacteria_ = 0.001; *R^2^_fungi_* = 0.05, *P*_fungi_ = 0.001). Globally, this structure was driven by the relative homogeneity of elevator samples in contrast to the strong dispersion effects observed across the other three environments (PERMDISP *P* = 0.001 for both domains). Interestingly, when compared directly to the tight clustering of their respective total community, the pathogenic subsets within elevators showed increased heterogeneity (**Figure 1g** and **1h**). This structural decoupling suggests that localized micro-environmental dynamics and stochastic human inputs may introduce greater compositional volatility within the pathogenic fraction than is found in the background core microbiome. –When all spatial groups were combined, clear seasonal partitioning emerged, particularly within the fungal domain (**Figure S4d**, PERMANOVA *R^2^_bacteria_* = 0.06, *P*_fungi_ = 0.001), where autumn and winter communities diverged distinctly from those in spring and summer. At the taxonomic level, this fungal macro-seasonal divergence was partially driven by the widespread enrichment of *Fusarium* across all spatial groups during the autumn and winter months (**Figure 2h**). Conversely, while the seasonal signal was less apparent (PERMANOVA *R^2^_bacteria_* = 0.06, *P*_bacteria_ = 0.001)) for the bacterial pathogenic community as a whole (**Figure S4c**), prominent temporal dynamics emerged at the taxon level within specific spatial contexts. This localized variation was highlighted by the pronounced enrichment of *Listeria* in aboveground and elevator spaces, contrasted by the preferential enrichment of *Helicobacter* in underground environments in spring (**Figure 2g**).

### Cross-Domain Ecological Networks Among Pathogenic Taxa Across Seasons

To determine if ecological interactions stabilized the pathogen distribution patterns, co-association networks integrating bacterial and fungal pathogen-containing genera were constructed for each season (**Figure 3**). Network robustness exhibited clear temporal variation (**Figure S5**). While natural connectivity declined year-round under targeted node removal, spring networks showed greater tolerance to the loss of highly connected nodes, and summer networks resisted the removal of bridging taxa. Strikingly, winter networks consistently displayed the lowest robustness and most rapid decline in connectivity. This indicates that pathogen interaction networks are substantially more vulnerable during the cold season, despite the concurrent peak in fungal pathogen abundance.

**Figure 3.**
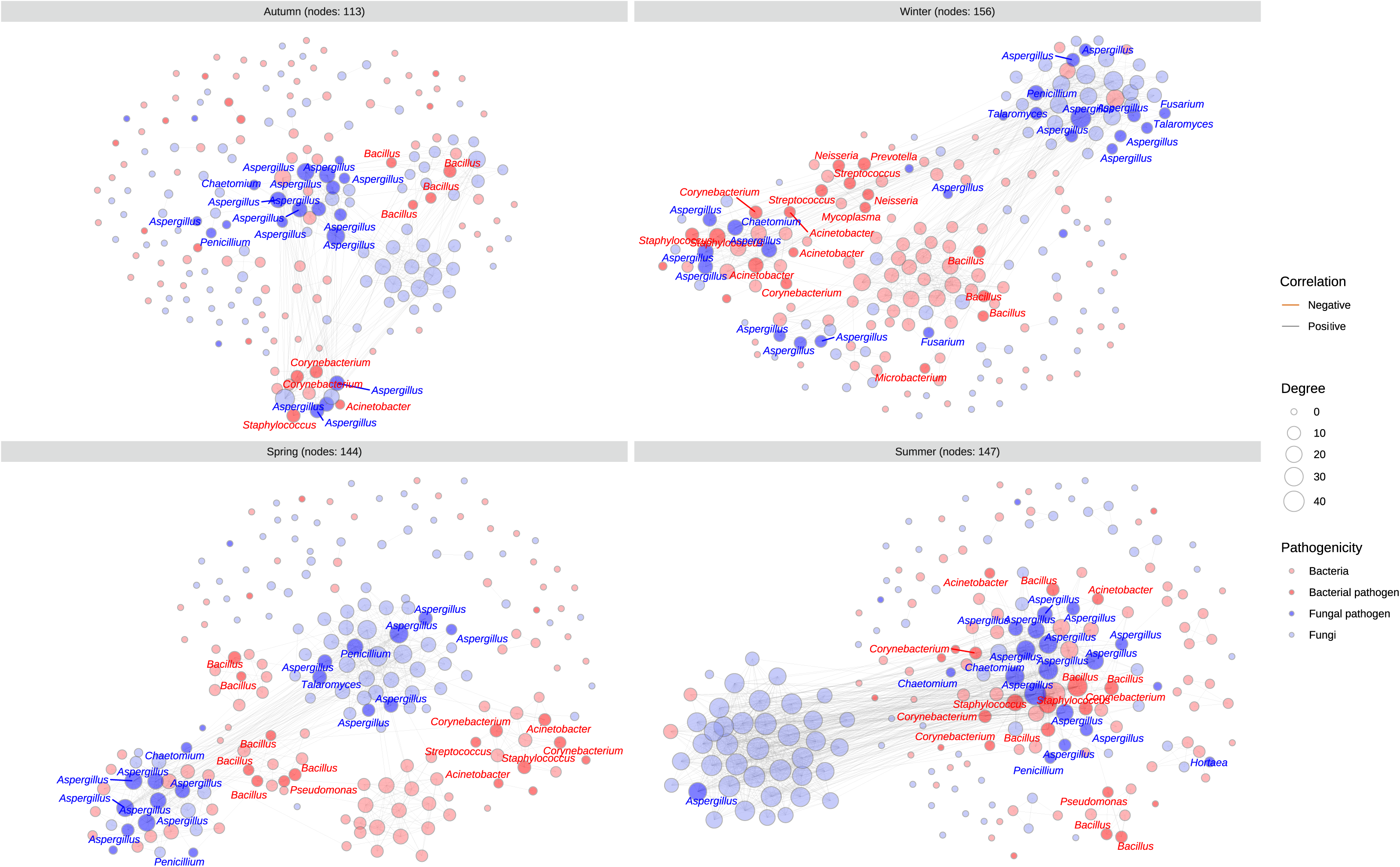
Cross-domain ecological networks of airborne microbial communities across seasons. Networks of inferred associations between bacterial and fungal ASVs were constructed using SPIEC-EASI. Only the top 200 most abundant bacterial and fungal ASVs were retained for network analysis. Each node represents an ASV and is colored by domain (bacteria or fungi) and pathogenicity status of the corresponding genus. Each edge indicates a significant association with an absolute coefficient greater than 0.6, and the color denotes the sign of the correlations. The network reveals extensive cross-domain interactions, suggesting potential co-occurrence and exclusion patterns among airborne microbes.

Co-association patterns revealed dense, predominantly positive interactions. The allergic fungal genus *Aspergillus*, which dominated across all the samples (**Figure 2h**), consistently occupied a keystone hub position across all four seasons, maintaining stable positive associations with another fungal genus *Chaetomium* throughout the year. Season-specific interaction restructuring was prominent: in autumn, *Aspergillus* associated positively with pathogenic bacterial and fungal genera, including *Corynebacterium*, *Staphylococcus*, and *Penicillium*. During winter, this hub expanded to encompass additional members, including *Talaromyces*, *Fusarium*, and *Acinetobacter*. A consistent positive association between fungal genera *Talaromyces* and *Penicillium* was also observed, reflecting coordinated dynamics among cold-favored fungi. Interdomain associations extended beyond *Aspergillus*; for example, *Chaetomium* correlated positively with the bacterial genera *Acinetobacter* and *Bacillus*. In spring, bacterial pathogen-containing genera formed tightly interconnected subclusters, in which *Staphylococcus* exhibited strong positive associations with *Acinetobacter*, *Bacillus*, *Corynebacterium*, and *Streptococcus*, indicating shared ecological niches or synchronous responses to environmental drivers. In summer, the network shifted toward a more bacteria-dominated structure with reduced intrafungal connectivity. In addition to the *Staphylococcus*-centered associations observed in spring, new positive correlations emerged between *Bacillus* and *Pseudomonas*, as well as between *Acinetobacter* and *Corynebacterium*, highlighting seasonal reorganization of bacterial co-occurrence patterns under warm-season conditions.

### Climatic Drivers of Pathogen Distribution at the Genus Level

Beyond the strong influence of spatial context, seasonal shifts in environmental conditions (i.e., relative humidity and temperature) exerted a secondary but crucial influence on community structure. At the community level, the richness of both bacterial and fungal communities exhibited a weak but significant negative correlation with relative humidity and temperature, indicating a general suppression under warmer, more humid conditions (**Figure S6**). However, the richness of both domains peaked at a relative humidity of 50-60% and a temperature of 15-17℃, a pattern more pronounced in fungi, suggesting a favored condition for the majority of microbes.

The responses of individual taxa, especially the pathogens, to the changes in environmental conditions were further assessed using MaAsLin3 (**Table S1**). Temperature exhibited a taxon-dependent influence. In terms of relative abundance, fungal pathogens such as *Talaromyces* and *Aspergillus* declined with increasing temperature, whereas a positive association was found in the bacterial pathogen *Escherichia* (**Figure 4a**). When considering only the presence/absence of taxa, several bacterial pathogens, including *Microbacterium*, *Mycobacterium*, and *Prevotella*, showed decreasing prevalence with increasing temperature (**Figure 4b**), indicating enhanced persistence under cooler conditions. A similar pattern was also found in fungal pathogens, including *Talaromyces*, *Sarocladium*, and *Chrysosporium* (**Figure 4c**).

**Figure 4.**
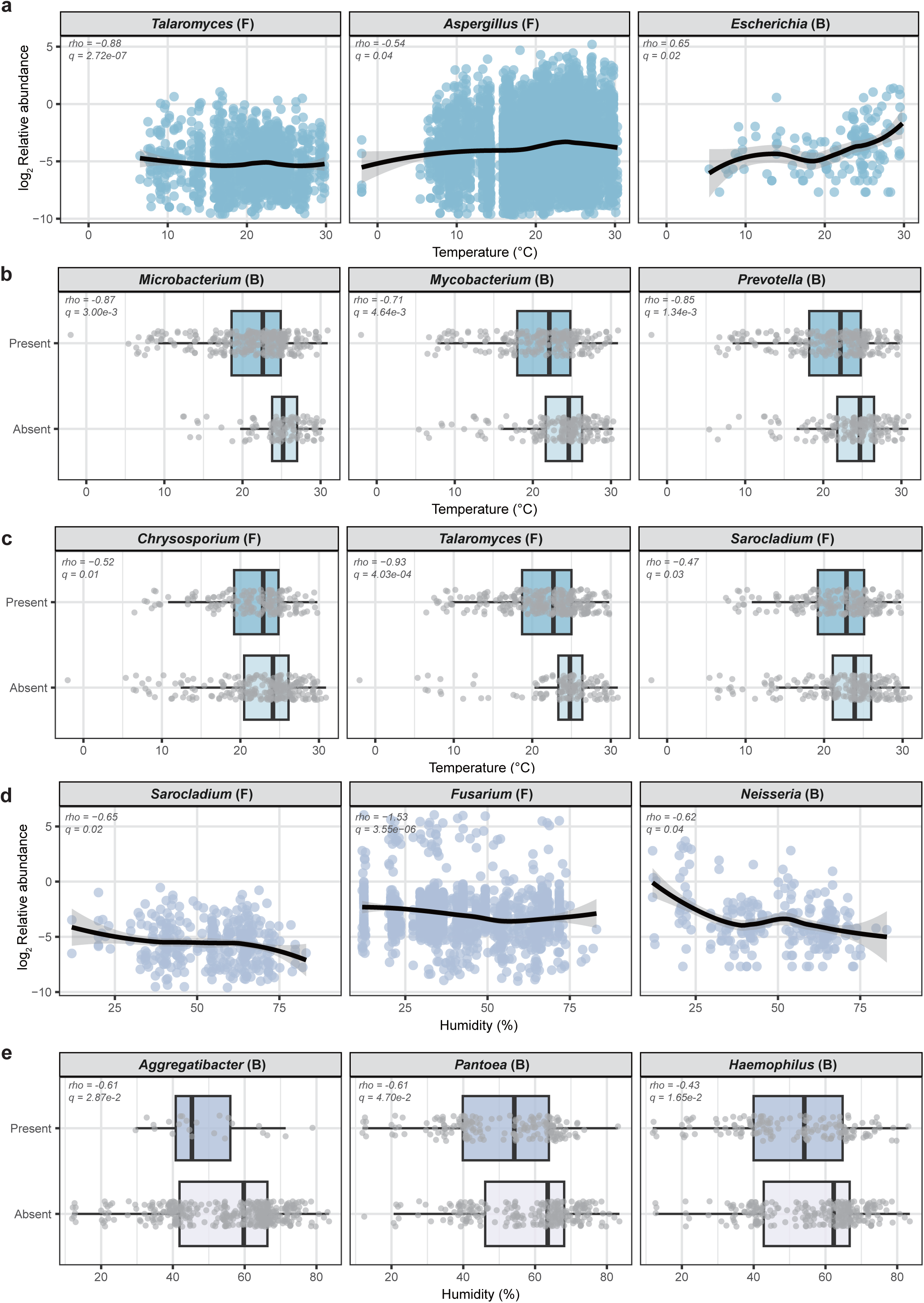
Environmental drivers of airborne bacterial and fungal pathogen dynamics. **a)** Correlations between temperature and the log₂-transformed relative abundance of pathogen-containing genera, including *Escherichia* (B), *Talaromyces* (F), and *Aspergillus* (F). **b)** Comparison of temperature between samples with and without detection of bacterial pathogen-containing genera (*Microbacterium*, *Mycobacterium*, and *Prevotella*). **c)** Comparison of temperature between samples with and without detection of fungal pathogen-containing genera (*Chrysosporium*, *Talaromyces*, and *Sarocladium*). **d)** Correlations between relative humidity and log₂-transformed relative abundance of bacterial and fungal pathogen-containing genera *Neisseria* (B), *Sarocladium* (F), and *Fusarium* (F). **e)** Comparison of relative humidity between samples with and without detection of bacterial pathogen-containing genera (*Aggregatibacter*, *Pantoea*, and *Haemophilus*). The labels B and F in the parentheses denote the respective microbial domain: B = Bacteria, F = Fungi.

Unlike temperature, relative humidity showed predominantly negative influence on both the prevalence and relative abundance of pathogen-containing genera. *Neisseria*, together with fungal pathogens such as *Sarocladium* and *Fusarium*, exhibited marked declines in relative abundance with increasing relative humidity (**Figure 4d**), suggesting a preference for drier environments consistent with their elevated winter representation (**Figures 2g and 2h**). Likewise, several bacterial pathogens, including *Aggregatibacter*, *Pantoea*, and *Haemophilus*, were more prevalent under less humid conditions (**Figure 4e**). Collectively, these results indicate that relative humidity and temperature act as key micro-environmental filters shaping pathogen-associated genera distributions in a highly taxon-specific manner.

### Seasonal Modulation of Bacterial Transcriptional Activity

Given the significant climatic filtering observed, we further examined the influence of environmental conditions on bacterial transcriptional activity by comparing rRNA-based (active) and rDNA-based (total) sequencing data from two climatically contrasting seasons: winter and the summer plum rain period. A weak but positive correlation between the relative abundance of bacterial members in the active and total community pools was observed in winter (Spearman’s ρ = 0.24, *P* < 0.001). In contrast, a similar trend was observed, but not statistically significant, during the plum rain period (Spearman’s ρ = 0.16, *P* = 0.1; **Figure 5a**).

**Figure 5.**
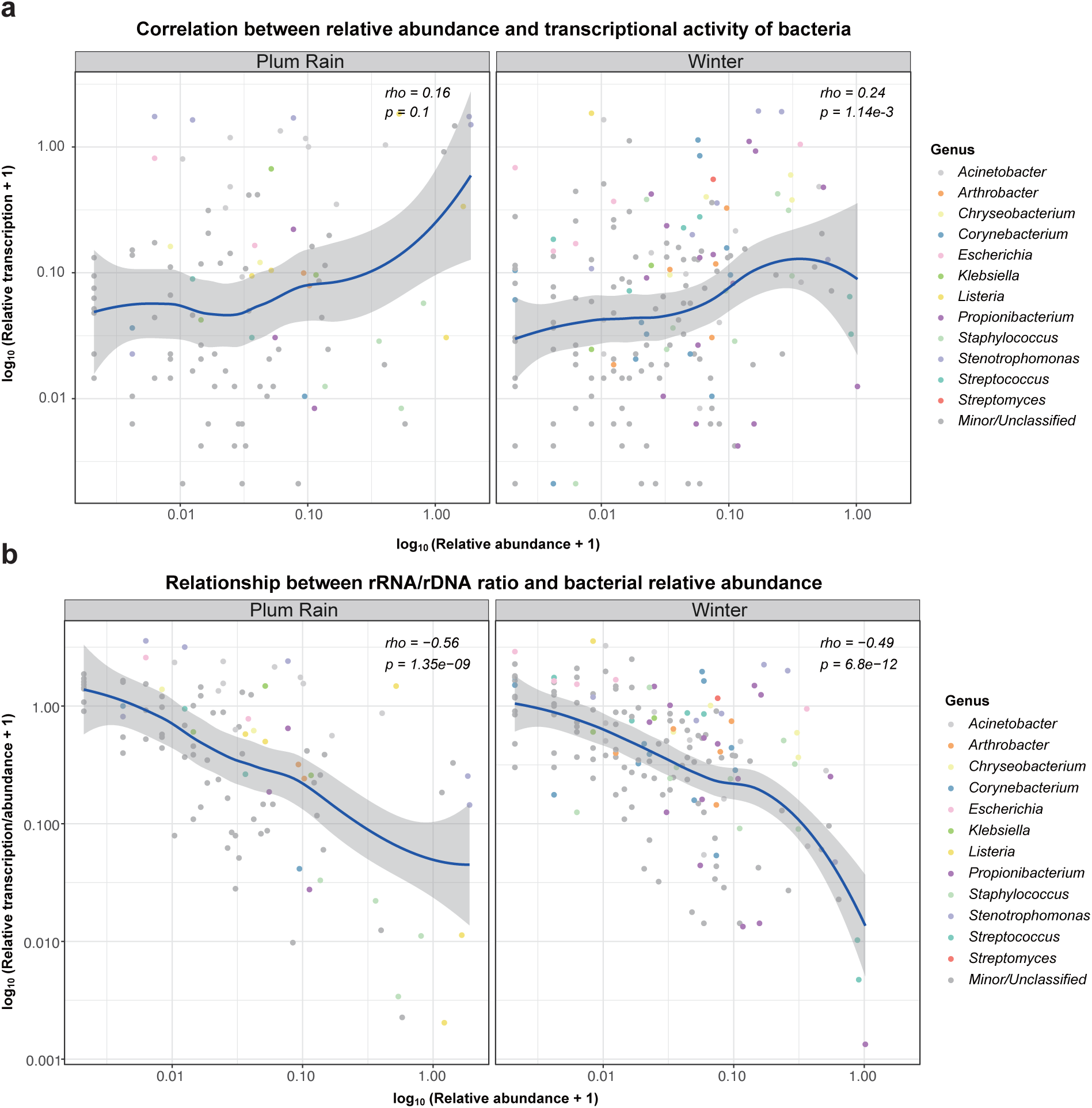
Transcriptional activity of airborne bacterial taxa in underground environments. **a)** Spearman correlations between relative abundance (rDNA) and relative transcriptional activity (rRNA) of bacterial ASVs in winter and plum rain summer season samples. **b)** Relationships between rRNA/rDNA ratio and relative abundance of bacterial ASVs across the same conditions. Each point represents an ASV detected in both rRNA and rDNA pools from the parallel samples. The blue line indicates the fitted linear regression trend. ASVs are taxonomically annotated at the genus level. A ratio >1 indicates putative metabolically active taxa, whereas values between 0 and 1 indicate low or inactive transcriptional states.

To rigorously investigate this relationship, we quantified the relative transcriptional activity of individual bacterial ASVs using the rRNA/rDNA ratio, where higher values are interpreted as a proxy for increased transcriptional activity. In both seasons, ASVs with higher relative abundances in the total community generally exhibited lower transcriptional activity (Spearman’s ρ_plum_rain_ = −0.56, ρ_winter_ = − 0.49, *P* < 0.001 for both seasons; **Figure 5b**), indicating that dominance in the total community does not necessarily translate to heightened metabolic vigor. Nonetheless, taxon-specific transcriptional dynamics were evident among co-detected pathogen-associated ASVs. Certain ASVs from *Acinetobacter*, *Escherichia*, and *Stenotrophomonas* maintained consistently high relative activity in both seasons. Pronounced seasonal contrasts also emerged: a *Klebsiella* ASV displayed enhanced transcriptional activity in summer, while ASVs from *Propionibacterium*, *Corynebacterium*, *Listeria*, and *Streptococcus* were metabolically more active in winter. These season-dependent bacterial transcriptional profiles complement the abundance dynamics and climatic correlations documented earlier, reinforcing the concept that environmental conditions not only reshape community composition but also actively modulate the metabolic state of persistent pathogens.

### Dispersal Dynamics and Cross-Spatial Transmission of Pathogen-associated Taxa

To determine the extent to which active dispersal drives these spatial and climatic patterns across different environments, we evaluated cross-spatial taxon sharing. A substantial core microbiota—comprising 53% of bacterial and 36% of fungal taxa—was ubiquitous across all four spatial settings (**Figure S7**). Crucially, this shared pool encompassed 35 pathogenic genera in each domain, highlighting the pervasive baseline distribution of clinically relevant microorganisms across complex BEs. We then quantified the transmission pattern of these pathogens between outdoor and indoor environments, and among distinct indoor spatial compartments using the Sloan NCM. Overall, both bacterial and fungal communities exhibited relatively limited migration rates, though bacterial taxa showed slightly higher dispersal potential than fungi (**Figure 6**). Among all modeled transmission routes, migration from aboveground and underground environments toward elevators consistently exhibited the highest rates for both domains. This identifies elevators as critical functional hubs for cross-spatial microbial exchange, a finding structurally congruent with their elevated diversity and tight bacterial–fungal coupling established in our previous analyses (**Figure 1k**).

**Figure 6.**
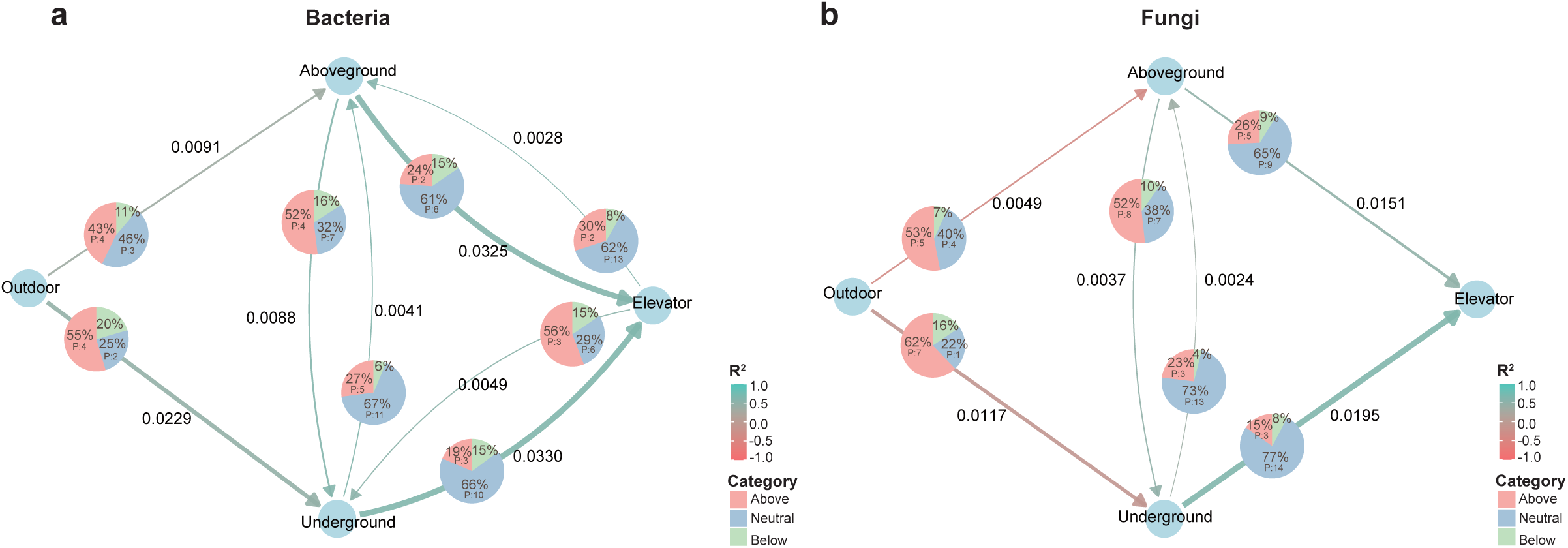
Inferred diffusion routes of airborne bacteria and fungi across four spatial environments. **a, b)** Inferred transmission pathways of bacterial **(a)** and fungal **(b)** communities among spatial environments within the buildings. Directional arrows indicate dispersal routes, with arrow thickness proportional to the coefficient of determination (R²), reflecting the strength of each pathway. The decimals adjacent to the arrows represent migration rates (*m*), where higher values denote greater dispersal potential. The embedded pie charts illustrate the proportions of ASVs distributed across the above-neutral, neutral, and below-neutral partitions; the label P within the above-neutral and neutral partitions indicates the specific counts of pathogen-containing genera unique to those environments.

Despite generally constrained macro-level dispersal, specific pathogen-associated taxa demonstrated potent migration capacities (**Table S2**). Within the fungal domain, *Aspergillus* and *Candida* consistently fell into the above-neutral category across all dispersal pathways, indicating active migration beyond neutral expectations. This strongly corroborates the ecological ubiquity and persistent abundance of *Aspergillus* observed throughout our study. *Alternaria* and *Talaromyces* also showed elevated dispersal signals across all four routes, while *Purpureocillium*, *Penicillium*, *Trichosporon*, and *Curvularia* exhibited above-neutral transmission in at least two pathways. Within the bacterial domain, *Acinetobacter*, *Streptococcus*, *Prevotella*, and *Clostridium* exhibited above-neutral dispersal across all four routes. This highlights their strong transmission capabilities, which directly support their broad spatial representation and central roles within the cross-domain interaction networks.

Ultimately, these findings establish that although community-level dispersal was generally constrained, specific pathogenic taxa exhibited disproportionately high dispersal potential, indicating that transmission processes operate heterogeneously across taxa. This active dissemination operates synergistically with spatial environmental filtering, climatic selection, and biotic interactions, forming an integrated mechanistic framework that governs the assembly, maintenance, and transmission of pathogen communities within architectural ecosystems.

## Discussion

BEs constitute the primary setting of human microbial exposure, yet the ecological mechanisms governing the distribution and persistence of pathogenic microorganisms within these systems remain incompletely understood. By integrating year-round observations across multiple spatial environments with community analyses and ecological modeling, this study revealed that pathogen-associated microbial communities were not randomly assembled but were structured by the combined effects of spatial filtering, seasonal forcing, biotic interactions, and dispersal. These processes operated in a coordinated manner, forming an interconnected ecological system that regulates pathogen occurrence, persistence, and transmission across space and time.

Among these processes, spatial filtering establishes the primary structure of pathogen-associated communities. Consistent with patterns observed in total microbial assemblages, both the diversity and composition of pathogenic subsets varied strongly across spatial environments, with enclosed indoor spaces exerting the strongest selective pressures. Notably, elevators exhibited a distinct ecological signature characterized by simultaneously elevated opportunistic pathogen diversity and limited community dispersion. This pattern reflects the interplay between continuous microbial input and deterministic environmental filtering. While frequent human-associated inputs introduce diverse taxa, restricted airflow and confined conditions create relatively stable conditions that impose persistent but monotonous selection, leading to a highly diversified microbial community co-occurring with reduced compositional variability (1). Similar dynamics have been reported in other enclosed, high-density environments, such as hospital wards, subway systems, and aircraft cabins, where architectural confinement and sustained human traffic collectively promote the accumulation and persistence of opportunistic pathogens, establishing these spaces as reservoirs with heightened pathogen transmission potential (23–25). Our results extend this framework by showing that elevators function not only as accumulation sites but also as selective environments that stabilize opportunistic pathogen assemblages, amplifying their ecological and epidemiological importance within buildings.

The strong spatial filtering resulted in the overall low migration rates of microorganisms across different spatial environments; nevertheless, several key taxa displayed effective cross-space transmission, possibly through human-mediated and airflow vectors (26). The spore-forming, opportunistic fungal pathogens, such as *Aspergillus* and *Candida*, were particularly prominent, reflecting their efficient dissemination and potential airborne transmission via small, resilient spores (27, 28). Similarly, several genera of bacterial opportunistic pathogens, including *Acinetobacter*, *Streptococcus*, *Prevotella*, and *Clostridium*, also exhibited broad distribution across multiple spatial compartments. Among these, human-associated taxa such as *Streptococcus* and *Prevotella* are likely redistributed through respiratory bioaerosols and skin shedding (29, 30). In contrast, desiccation-tolerant environmental genera such as *Acinetobacter* and *Clostridium* may persist on inanimate surfaces, such as ventilation filters, under drier conditions and be secondary aerosolized and further dispersed via disturbance-driven dust resuspension (31, 32).

Superimposed on this spatial baseline, seasonal climatic variation further modulates pathogen dynamics, but with markedly different effects on bacterial and fungal communities. Opportunistic fungal pathogens exhibited pronounced seasonal shifts, with elevated relative abundance across four spatial environments during autumn and winter. This pattern does not necessarily reflect optimal fungal growth conditions, as most indoor fungi thrive at moderate temperatures (typically 20-30°C) and high relative humidity (> 70%), as previously reported (33). Instead, this phenomenon may result from a combination of increased spore persistence, reduced microbial competition, and enhanced aerosolization under low-humidity conditions rather than *in situ* growth dynamics (33, 34). For instance, the spore-producing fungal genus *Fusarium* generally prefers moist environments for active growth (35); thus, its increased relative abundance during autumn and winter likely reflects enhanced environmental loading from plant-derived sources (36), combined with improved aerosolization under drier atmospheric conditions (37), thereby increasing suspension time and transport efficiency. Concurrently, the reduced activity of other competing taxa during colder periods may create ecological opportunities that allow *Fusarium* to persist, even if its intrinsic growth rate is suboptimal.

The temporal dynamics of opportunistic bacterial pathogens appeared to be governed by taxon-specific physiological strategies rather than uniform seasonal responses. Although *Listeria* exhibited lower relative abundance during winter, its transcriptional activity was paradoxically higher than in summer. Rather than contradicting the capacity of bacterial pathogens to persist under cold stress, this pattern is consistent with the well-established cold-adaptation strategy of *Listeria* species such as *L. monocytogenes*, which actively upregulates cold shock proteins and stress-associated sigma factors at low temperatures to maintain cellular viability (38). The elevated transcriptional activity observed in winter, therefore, likely reflects active stress adaptation rather than metabolic dormancy. With the transition into spring, the relaxation of cold-induced stress, together with moderate increases in temperature and relative humidity, may shift cellular resource allocation from stress tolerance toward growth and replication, thereby promoting population expansion and the marked increase in relative abundance observed during this period. Simultaneously, the seasonal decline in fungal dominance may partially alleviate interkingdom competitive constraints, further creating ecological space favorable for the proliferation of opportunistic bacterial pathogens. These asynchronous seasonal dynamics between two domains indicate that pathogen risk does not diminish uniformly throughout the year but instead shifts in composition, with opportunistic fungal pathogens dominating under colder conditions and bacterial counterparts becoming more prominent during transitional and warmer periods.

Beyond spatial and environmental constraints, internal biotic interactions play a critical role in shaping pathogen persistence. Cross-domain network analyses revealed that pathogenic taxa are embedded within densely connected and predominantly positive interaction networks. These consistent robust co-occurrence patterns may reflect shared ecological preferences, localized environmental filtering, or potential mutually beneficial associations. Notably, the allergic fungal genus *Aspergillus* persistently occupied a central hub position across all seasons. While its keystone role may be attributed to a combined strategy of ecological versatility, desiccation tolerance, and continuous dispersal via airborne spores (7, 28), its network centrality suggests it may act as a physical or structural anchor for other taxa. In generally dry BEs, dense fungal structures and spore accumulations can alter surface topography, creating micro-niches that retain trace moisture and organic matter, thereby facilitating the attachment and survival of other fungal and bacterial partners (39–41). This combination of high physical tenacity and localized niche stabilization likely drives the strong, positive cross-domain coupling centered around *Aspergillus*, ultimately enhancing the collective persistence of pathogenic assemblages within the indoor biome.

Crucially, the seasonal variation in network topology indicates that pathogen interactions are dynamically reorganized rather than statically maintained. Although our structural analyses showed that winter networks were more vulnerable to targeted node removal, they did not collapse; rather, they shifted toward a heavily fungus-dominated configuration with sustained cross-domain associations. This structural reorganization is likely driven by the seasonal enrichment of the dominating genus *Fusarium* and the less common genus *Talaromyces*. In contrast, the dominance of bacterial pathogens in the summer network, including *Bacillus*, *Staphylococcus*, and *Corynebacterium*, was accompanied by reduced intrafungal connectivity. These seasonal transitions suggest that pathogen interaction networks are shaped by distinct ecological strategies across climatic conditions: winter communities appear to be maintained largely through the persistence and dispersal of fungal spores, whereas summer communities are characterized by enhanced bacterial activity and intensified bacterial co-association patterns. Collectively, these findings demonstrate that seasonal dynamics reshape not only community composition, but also the fundamental balance and interaction architecture of pathogen networks.

Taken together, our study supports a unified framework in which pathogen dynamics in BEs emerge from the coupled effects of spatial filtering, climatic modulation, and biotic interactions. Within this system, active dispersal facilitates the initial colonization of dominating pathogenic taxa, while subsequent cross- and intra-domain biotic interactions promote their long-term surface persistence. Elevators emerge as critical nodes within this framework; the stable environmental conditions they provide, coupled with relatively high inbound migration rates, position them as primary reservoirs for microbial pathogens. Because these ecological processes are intrinsically linked, they jointly govern pathogen persistence and redistribution across the building. This perspective has clear, actionable implications for future intervention strategies. Approaches targeting a single factor, such as uniform disinfection or static environmental control, are less effective in disrupting this coordinated system. More effective management should prioritize structurally critical nodes, such as the ventilation interfaces within elevators, while dynamically accounting for seasonal shifts in dominant pathogen groups and the stabilizing roles of keystone taxa. Looking forward, as climate change reshapes indoor temperature and humidity regimes, progress toward predictive pathogen control will depend heavily on integrating microbial activity profiling with high-resolution micro-environmental monitoring and airflow-informed architectural modeling.

These conclusions should be interpreted in light of several limitations. Amplicon-based sequencing optimizes broad community profiling but restricts taxonomic resolution to the genus level, limiting the precise identification of pathogenic species and their virulence traits. Furthermore, this pathogenic genus-level assignment may overestimate pathogen prevalence by including nonpathogenic members within broadly defined taxa. Transcriptional activity was assessed primarily for bacterial communities and across two seasons, providing an initial framework that may not fully capture the full extent of functional dynamics, particularly for fungi. In addition, it should be noted that rRNA-based metrics provide a valuable proxy for estimating transcriptional activity, though they may be influenced by variation in ribosomal copy number and physiological states. Finally, neutral modeling and co-occurrence networks can infer potential dispersal and interactions, but they do not establish strict causality. Future work integrating metagenomics, metatranscriptomics, and quantitative airflow modeling will be essential to further elucidate these mechanisms and improve the prediction of pathogen dynamics in BEs.

## Acknowledgements

We thank the participants who contributed to the sample collection.

## Authors’ contributions

S.Q. and S.Z. performed bioinformatic analysis, data analysis, and interpretation, and wrote the manuscript. Y.H., M.S., Z.X., S.B., and Y.S. performed sampling and genomic DNA extraction. L.S. performed total RNA extraction and reverse transcription. X.T. conceived the study, supervised the research, and wrote the manuscript.

## Data, metadata, and code availability

The raw DNA and RNA sequencing data for bacterial and fungal communities have been deposited in the NCBI Sequence Read Archive (SRA) database under the BioProject accession numbers PRJNA1268906 and PRJNA1466084, respectively. The metadata and in-house scripts for data analysis are available at https://github.com/Shijiao7/XJTLU_Air_Microbiome

## Supplementary Materials

**Text S1. Supplementary method file.**

**Table S1. MaAsLin3 results for associations between microbial taxa and metadata covariates.**

**Table S2. Sloan neutral community model quantifying transmission of pathogen-containing genera.**

**Figure S1. Overview of the spatiotemporal sampling strategy and environmental microclimates during the study period. (a)** Chronological timeline of the sampling campaigns, illustrating the specific data collection points mapped across four continuous seasons. **(b)** Schematic floor plan detailing the spatial distribution of sampling sites within the representative underground environment. Distinct functional zones (e.g., elevators, offices, canteens) and specific sampling nodes are marked to capture the spatial configuration of the built environment. **(c)** Variation in mean relative humidity and temperature fluctuations were continuously monitored across the four defined spatial types and sampling dates.

**Figure S2. Seasonal dynamics of microbial richness inferred using generalized additive models (GAMs). a, b)** Observed log-transformed richness of bacterial **(a)** and fungal **(b)** communities over time. Solid lines indicate GAM fits, with shaded areas representing 95% confidence intervals. **c, d)** Estimated seasonal trends derived from cyclic splines, showing annual periodicity while controlling for long-term trends.

**Figure S3. Boxplot of Procrustes residuals from the analysis of bacterial and fungal community dissimilarity.**

**Figure S4. Diversity analyses for bacterial and fungal opportunistic pathogen-containing genera in different seasons. a, b)** Shannon diversity of bacterial **(a)** and fungal **(b)** pathogen-associated communities across four seasons. **c, d)** Principal coordinates analysis (PCoA) of the pathogen-associated communities in four seasons, using Bray–Curtis dissimilarity for both domains.

**Figure S5. Seasonal variation in cross-domain network stability. a, b)** Network robustness in each season was assessed by natural connectivity under targeted node removal. **a)** Betweenness-based attack: nodes were sequentially removed in descending order of betweenness centrality in the networks. **b)** Degree-based attack: nodes were removed in descending order of node degree in the networks. Lines represent the change in natural connectivity as a function of the fraction of nodes removed, with higher values indicating greater network stability.

**Figure S6. Associations between microbial richness and environmental variables. a, b)** Spearman correlations between microbial richness and environmental variables, including temperature and relative humidity, for bacterial **(a)** and fungal **(b)** communities. Correlation coefficients (ρ) and significance levels are shown for each relationship.

**Figure S7. Shared and unique microbial genera across four spatial environments. a, b)** Venn diagrams showing the overlap and uniqueness of bacterial **(a)** and fungal **(b)** genera among the four spatial groups. Numbers indicate the count of genera shared among or unique to each spatial environment.

